# A pan-cancer interrogation of intronic polyadenylation and its association with cancer characteristics

**DOI:** 10.1101/2022.11.14.516018

**Authors:** Liang Liu, Peiqing Sun, Wei Zhang

## Abstract

mRNA cleavage and polyadenylation is an integral 2-step process in the generation of protein-encoding mRNA or noncoding transcripts. More than 60% of human genes have multiple polyadenylation sites either in the 3’ untranslated region (3’UTR-APA) or in the intronic/exonic region, resulting in expression of isoforms with alternative polyadenylation (APA) under different physiologic conditions. The 3’UTR-APAs have been extensively studied, but the biology of intronic polyadenylations (IPA) remain largely unexplored. Here we characterized the IPA profiles of 9,679 patient samples across 32 cancer types from the Cancer Genome Atlas (TCGA) cohort. Overall, we identified 22,260 detectable IPA sites; 9,014 (40.5%) occurred in all 32 cancer types and 11,676 (52.5%) occurred in 2 to 31 cancer types. By comparing tumors and their paired normal tissues, we identified 180 to 4,645 dysregulated IPAs in 132 to 2,249 genes in each of 690 patient tumors from 22 cancer types that showed consistent patterns within individual cancer types. Furthermore, across all cancer types, IPA isoforms and their gene regulation showed consistent pan-cancer patterns, and cancer types with similar histologic features were clustered at higher levels of hierarchy. We selected 2,741 genes that were consistently regulated by IPAs across cancer types, including 1,834 pan-cancer tumor-enriched and 907 tumor-depleted IPA genes. Pan-cancer tumor-enriched IPA genes were amply represented in the functional pathways such as cilium assembly and DNA damage repair. Expression of IPA isoforms in DNA damage repair genes was associated with tumor mutation burdens. Expression of IPA isoforms of tumor-enriched IPA genes was also associated with patient characteristics (e.g., sex, race, cancer stages, and subtypes) in cancer-specific and feature-specific manners. Importantly, IPA isoform expression for some genes could be a more accurate prognostic marker than gene expression (summary of all possible isoforms). In summary, our study reveals the roles and the clinical relevance of tumor-associated IPAs in cancer.

## INTRODUCTION

The process of mRNA maturation includes multiple steps. mRNA cleavage and polyadenylation (CPA) are key final steps that begin with cleavage of the 3′ end of the precursor mRNA followed by the sequential addition of adenosine to form a poly(A) tail (1,2). Over half of all human genes harbor multiple polyadenylation sites (PAS) that generate alternative cleavage and polyadenylation (APA) isoforms (3-5). Dynamic APA occurs in numerous physiologic processes such as cell development, differentiation, proliferation, and reprogramming (6-8), and in human diseases such as viral infection (9) and cancers (10-14). Cleavage sites lie between upstream and downstream mRNA sequence core elements. The core binding factors (i.e. CPSF, CstF, CFIm, and CFIIm) – and other proteins such as polyadenylation polymerase, gene/tissue-specific RNA binding proteins, and other trans-acting factors (15,16) – directly bind or associate through complexes to mRNA core regulatory cis-elements (e.g., AAUAAA and other auxiliary sequences such as U/GU-rich downstream elements (1)). Thus, genetic aberrations in these cis-elements lead to global APA reprogramming (17,18). Alternative splicing (and splicing factors (19-23)) and chromatin remodeling (e.g. DNA methylation (24) and m6A methylation (25,26)) also function in global APA process.

APA primarily occurs in the 3’-most exons of genes (3’UTR-APA); by changing the position of PAS, one gene produces multiple transcripts with either shorter or longer 3’UTRs, resulting in gain or loss of important cis-regulatory elements, such as miRNA binding sites (4). Intronic polyadenylation (IPA) of genes causes a stop of normal transcription, generating truncated mRNA isoforms and dysfunctional proteins or non-coding RNAs (ncRNAs) that do not produce proteins at all (1). IPA is involved in inactivation of DNA damage repair genes (27,28) and tumor suppressors (29) in cancer, and in transcript diversity in immune cells (30). Nevertheless, the clinical relevance of IPAs is poorly understood.

Various experimental approaches have been developed to evaluate PAS in a genome-wide manner (31-34). Applications of these techniques across normal and abnormal tissues enable the study of regulatory mechanisms and functional consequences of APA status, and to advance our knowledge of this post-transcriptional process under various biological conditions (6-11). However, these approaches are relatively laborious and/or costly in contrast to conventional RNAseq, which has been widely applied to elucidate the transcriptomics landcape of large-scale sample cohorts, such as the Cancer Genome Atlas (TCGA) (35) and the Genotype-Tissue Expression (GTEx) (36) cohorts. In this study, we employed a bioinformatics approach to quantify IPAs from conventional RNAseq data of samples from TCGA and illustrated systematic patterns of IPA isoforms, resultant impaired gene transcription, and their clinical relevance.

## MATERIALS AND METHODS

### Quantification of IPA isoform expression

PolyA_DB3 (37) was downloaded from https://exon.apps.wistar.org/PolyA_DB/v3/misc/download.php. PolyA_DB3 (37) consists of more than 290,000 PAS from 24 experiments, including ∼32,000 PAS mapped in the intronic regions of ∼9,000 genes (IPA sites). We applied APAlyzer (38) to quantitate IPA usage by defining genomic regions according to IPA site(s) within one gene and counting RNA-seq reads mapped into each region (**Fig 1B**). The gene coordinates were determined using the human genome reference (GRCh37 version). IPA-specific regions were defined as those spanning from the upstream 5′ splice site (5’ss) of the intron to each IPA site.

**Figure 1.**
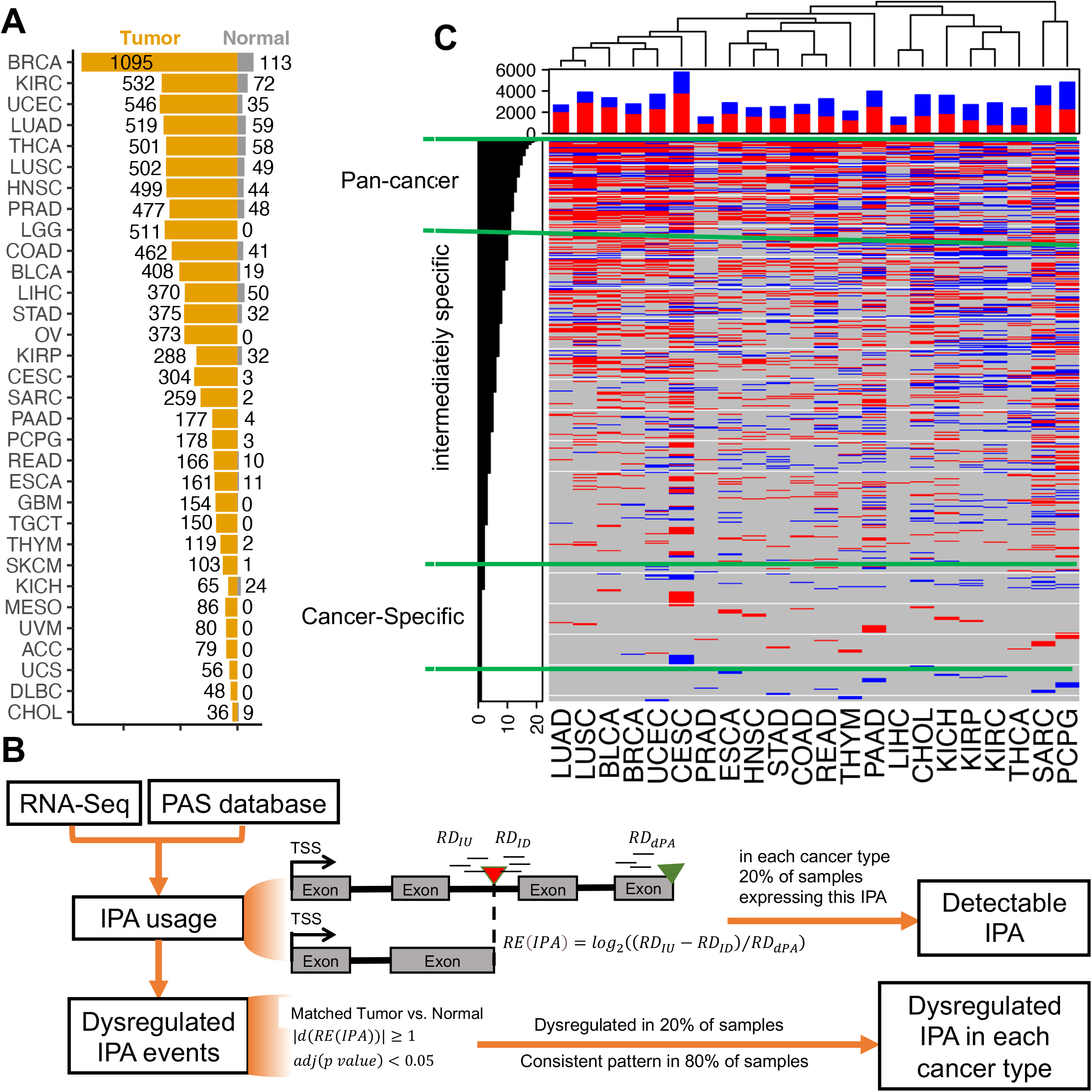
IPA landscape in the TCGA cohort. (A) Data source for the 33 cancer types in this study. Bar charts describe numbers of tumor (yellow) and normal samples (grey) with available RNAseq for each cancer type (cancer abbreviations are shown in **Table S1A**). (B) Workflow and criteria for IPA quantification using RNAseq data. RE: relative expression; dPA: distal PA; IU: IPA upstream; ID; IPA downstream; RD: read density. (C) IPAs (row) with increased (red) or decreased (blue) usage in each tumor type (column). The upper histogram shows the numbers of IPAs with increased (red) or decreased (blue) usages in each tumor type. The side histograms show the numbers of cancer types with dysregulated IPAss. A hierarchical clustering method was used. Pan-cancer IPA sites (n = 2,011): consistently dysregulated in 11 cancer types, intermediately specific IPAs (n = 7,570): consistently dysregulated in 2-10 cancer types, and cancer specific IPAs (n = 2,394): dysregulated in only 1 cancer type.

We downloaded RNA-seq BAM files from the TCGA data portal (https://portal.gdc.cancer.gov/) (35). RNA-seq reads exactly mapped to the IPA-specific region but not spanning 5’ss or IPA were considered IPA supporting reads (*NR*(*IPA*)) and used to calculate the reads density (RD) of IPA (*RD*_*IU*_). Reads exactly mapped to the 3’UTR region but not spanning the distal PAS (dPA; determining the end of 3’UTR) were considered as supporting reads of dPA (*NR*(*dPA*)) and used to calculate the reads density of full-length mRNA (*RD*_*dPA*_ ). The relative expression of one IPA was calculated as *RE*(*IPA*) = *log*_2_((*RD*_*IU*_ − *RD*_*ID*_)/*RD*_*dPA*_) ( *RD*_*ID*_ is the read density of the intronic downstream region of the IPA site to the 3’ss of the next exon).

### Identification of aberrent IPAs and IPA-regulated genes

We identified IPA isoforms that were differentially expressed between tumor and matched normal samples in the 22 cancer types where at least 2 matched tumor and normal sample were available (**Fig 1A**). A significantly dysregulated IPA was determined by its expression difference ( |*d*(*RE*(*IPA*))| ≥ 1 ) with an adjusted *p* < 0.05 from the Fisher’s exact test using the reads numbers in IPA-specific (*NR*(*IPA*)) and 3’UTR regions (*NR*(*dPA*)) in the two conditions (**Fig 1B**).

In each cancer type, an IPA was considered as consistently having higher or lower usage if it was dysregulated in ≥ 20% tumor samples and shows a consistent increasing or decreasing pattern in ≥ 80% of these samples (**Fig 1B**).

At gene level, a gene was considered to be transcriptionally truncated (**tumor-enriched IPA gene** that expresses the truncated isoform) if any IPA of this gene has a higher usage in tumors; a gene was considered as a **tumor-depleted IPA gene** that expresses the full-length isoform using polyadenylation sites in 3’UTR if any IPA of this gene has a lower usage but none has a higher usage (**Fig 2A**).

**Figure 2:**
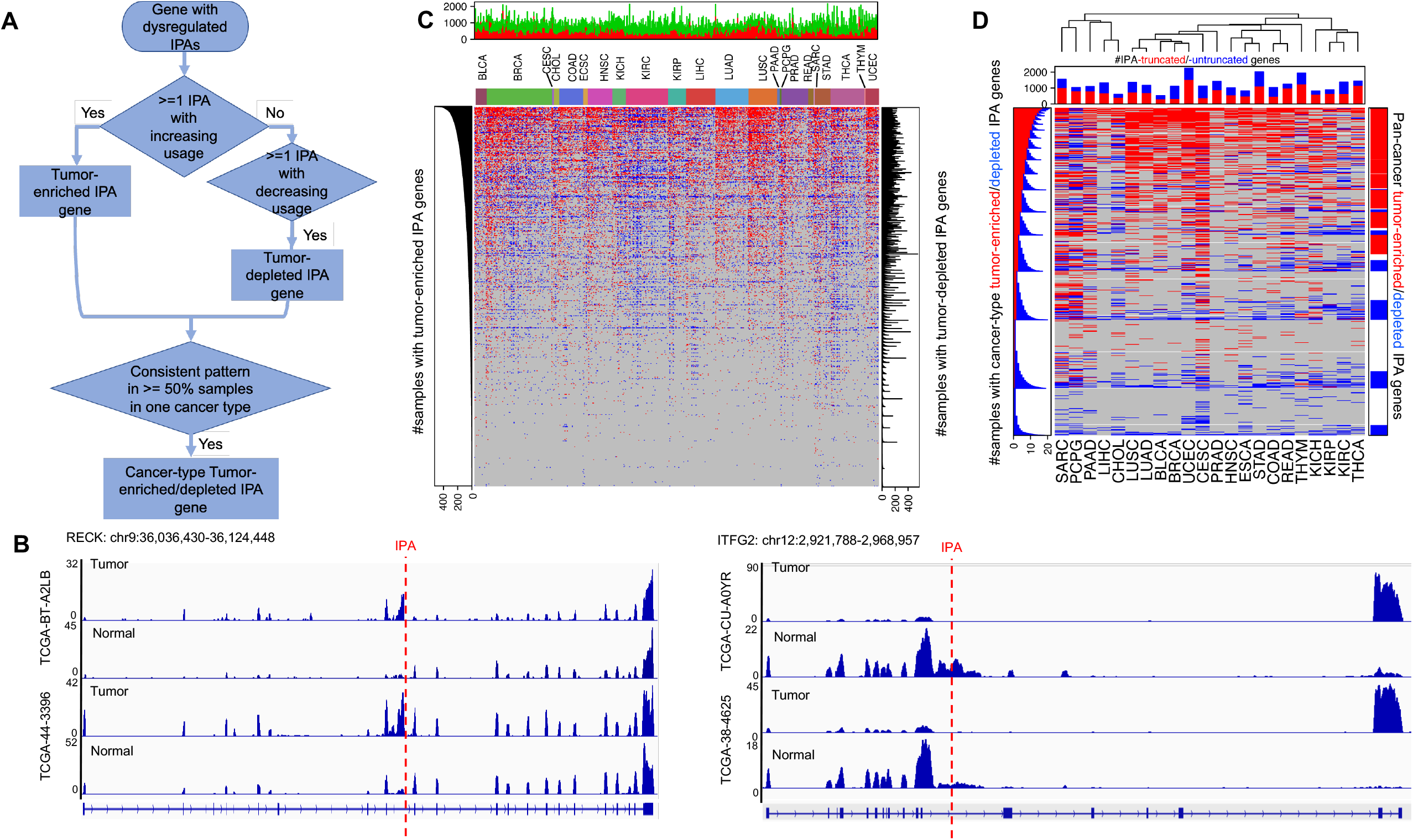
IPA-regulated genes across TCGA tumor types. (A) Strategy to determine if one gene was considered as a tumor-enriched or depleted IPA gene in each cancer type. (B) Example of tumor-enriched (RECK) and depleted (ITFG2) IPA genes in tumors compared to the matched normal samples. (C) The central heatmap shows tumor-enriched (red) or depleted (blue) IPA genes in each tumor (columns). The upper histogram shows the number of genes per tumor. The side histograms show the numbers of tumors with tumor-enriched (left) or depleted IPA (right) genes. The central heatmap shows cancer-type tumor-enriched or depleted IPA genes in each cancer type. The upper histogram shows the numbers of tumor-enriched (red) and depleted (blue) IPA genes in each cancer type. The left histograms show the numbers of cancer types with cancer-type tumor-enriched (red) and depleted (blue) IPA genes. The right histograms show whether the gene is selected as a pan-cancer tumor-enriched (red) and -depleted (blue) IPA gene. A hierarchical clustering method was used.

In each cancer type, we selected the cancer-type tumor-enriched IPA gene if this gene was determined as tumor-enriched IPA gene in more tumor samples (≥ 50% and at least two), and the tumor-depleted IPA gene if one was determined as cancer-type tumor-depleted IPA gene in more samples (≥ 50% and at least two) (**Fig 2A**).

Finally, we selected 1,834 and 907 pan-cancer tumor-enriched and -depleted IPA genes that were determined as tumor-enriched and -depleted IPA genes in at least half of the cancer types (**Fig 2D**).

### Coding potentials of IPA isoforms

To analyze coding potentials of IPA isoforms from their transcript sequences, we input the isoform transcript sequences (obtained based on gene and IPA coordinates and genome sequences from Gencode v22) to CPAT (39) to predict coding probability. We considered an isoform as a protein-coding mRNA if its coding probability score was greater than 0.364, and as a non-coding RNA if its coding probability score was < 0.364. Protein sequences were predicted using EMBOSS Transeq (40) available at the USCS genome browser.

### Functional enrichment analyses

Functional (gene ontology and hallmark) enrichment for selected gene sets was performed using the enrichR (41) package in R. Three scores were calculated: A p-value computed using Fisher’s exact test to determine significant overlap between other publicly available datasets, a z-score computed by assessing the deviation from the expected rank, and a combination of the adjusted p-value and the z-score. Enrichment of individual gene sets was considered significant if the adjusted p-value was < 0.05, unless stated otherwise.

### Truncating mutations, DNA damage repair gene, tumor suppressor gene and Tumor mutation burden

Somatic mutations were downloaded from the TCGA data portal (https://portal.gdc.cancer.gov/) (35). Nonsense mutations, frame-shift mutations, and splice-site mutations were considered as truncating mutations (29). DNA damage repair gene set were obtained from Gene Ontology database included in the enrichR (41) package. Tumor suppressor genes (42) were obtained from previous reports.

The patient tumor mutation burden matrix was downloaded from cBioportal (43,44) (https://www.cbioportal.org). 8,039 samples of the 22 cancer types were included in analysis. In each cancer type, patients were categorized into two groups (≥ or < 90%) based on expression levels of individual IPA isoform. Association between IPA isoform expression and tumor mutation burden was evaluated with two-sided Mann-Whitney test for each IPA isoform in each cancer type.

### Tumor immune infiltration

The relative proportions of 22 immune cell types within the leukocyte compartment of individual tumor samples were obtained from previous reports (45) using CIBERSORT (46). This tool uses a set of 22 immune cell reference profiles to derive a base (signature) matrix that can be applied to mixed samples to determine relative proportions of immune cells. Then these proportions were multiplied by the leukocyte fraction to yield corresponding estimates in terms of overall fraction in tissue. Values were aggregated in various combinations to estimate the abundance of more comprehensive cellular classes, such as lymphocytes, macrophages, and CD4 T cells (46). 8,039 samples of the 22 cancer types were included in analysis. In each cancer type, patients were categorized into two groups (≥ or < 90%) based on expression levels of individual IPA isoform. Proportion difference of each immune cell type was accessed with two-sided Mann-Whitney test.

### Determination of upstream regulatory factors for IPAs and effects of IPA-ncRNAs on signaling pathways

Polyadenylation and splicing factors were obtained from Gene Ontology database included in the enrichR (41) package. The mRNA expression matrix from the TCGA data portal (https://portal.gdc.cancer.gov/) (35). 8,039 samples of the 22 cancer types were included in analysis. In each cancer type, we used partial correlation (pcor in R) to evaluate correlations between individual IPA isoforms and regulator factors, including 42 polyadenylation factors and 262 splicing factors. Factors with |Rs| ≥ 0.3 and a false discovery rate [FDR] <0.05 were considered as statistically significant regulators. We further defined putative master regulators as those significantly correlated to ≥ 20% of individual IPAs in a specific cancer type. To facilate calculations, only consistently dysregulated IPAs in each cancer types (occurring in ≥ 20% of tumor samples compared to matched normal samples) were included.

Ten Cancer signaling pathways including p53, PI3K, Myc, RTK/RAS, cell cycle, Wnt, TGF beta, Nrf2, Notch, and Hippo were collected by previous reports (47). We used Spearman correlation to calculate correlations between the IPA-ncRNAs and gene members of each pathway. We defined the significant correlation with absolute value of Spearman correlation |Rs| > 0.3 and FDR < 0.05. Only IPA-ncRNAs produced by pan-cancer tumor-enriched IPAs were included.

### Clinically relevant IPAs

We obtained demographic and clinical information associated with tumor samples, including tumor subtypes, disease stage, patient race (black versus white), patient sex, and overall survival time from TCGA pan-cancer papers or the GDC data portal (https://gdc-portal.nci.nih.gov/). 8,039 samples of the 22 cancer types were included in analysis. In each cancer type, we used Mann-Whitney tests for two groups and analyses of variance (ANOVA) for multiple groups to assess differences among patient cancer subtypes and stage (r <0.05). All statistical tests were two-sided.

For survival analysis in each cancer type, patients into two groups (≥ or < 90%) based on levels of IPA expression or gene expression (sum of all isoforms e.g. those from alternative polyadenylation and splicing) of each gene. The mRNA expression matrix was download from the TCGA data portal (https://portal.gdc.cancer.gov/) (35). We used univariate Cox models or log-rank tests to assess whether IPA isoform or gene expression was associated with the overall survival times of cancer patients and considered p < 0.05 as significant.

## RESULTS

### Broadly and recurrently aberrant IPAs across tumor types

We collected RNAseq data from 9,679 tumor tissues across 32 cancer types from TCGA (35) (**Fig 1A; Table S1A**) and employed the APAlyzer (38) to measure usage of ∼32,000 IPAs within ∼9,000 genes, obtained from the PolyA_DB3 database (37). RNAseq reads were mapped to specific regions regarding polyadenylation sites (**see Methods**). We required one IPA to have at least 5 supporting RNAseq reads (*NR*(*IPA*) ≥ 5) and relative expression *RE*(*IPA*) ≥ *log*_2_(0.05) to be considered as an expressed IPA for inclusion in downstream analyses (**Fig 1B**). Across all tumor samples, we observed 22,260 IPA sites, with esophageal carcinoma (ESCA) having the highest and uveal melanoma the lowest median IPA burden (13,781 IPAs versus 10,558 IPAs, respectively, per sample) (**Fig S1A**). Overall, tumor samples had higher IPA burdens than normal samples except for a few cancer types, such as pan-kidney cancers (48) (kidney renal papillary cell carcinoma [KIRC], renal clear cell carcinoma [KIRP] and Chromophobe [KICH]) (**Fig S1B)**.

Next, we identified IPAs with differential usages between matched tumor and normal samples (n=690) in each of the 22 tumor types (having at least 2 matched samples) (**Table S1B**). We defined a significantly dysregulated IPA based on its relative expression difference (|*d*(*RE*(*IPA*))| ≥ 1) and adjusted *p* < 0.05 from Fisher’s exact test (**Fig 1B; see Methods**). The number of differential IPAs varies between 180 to 4,645 in 132 to 2,249 genes among the 690 tumor samples (**Fig S1C**). In addition, most of the IPAs are consistently dysregulated in tumors compared to matched normal samples within individual cancer types (**Fig S1D**).

We selected IPAs with consistent patterns in each cancer type for analysis (**see Methods**). Across all cancer types, cervical squamous cell carcinoma and endocervical adenocarcinoma CHOL cholangiocarcinoma (CESC) has the highest number of dysregulated IPAs, followed by pheochromocytoma and paraganglioma (PCPG); the fewest dysregulated IPAs are in samples of liver hepatocellular carcinoma (LIHC) (**Fig S1E; Table S1B**). Most cancer types show strikingly elevated IPA usage, such as lung adenocarcinoma (LUAD) and squamous carcinoma (LUSC). However, five cancer types show opposite trends, including KIRC, thyroid carcinoma (THCA), cholangiocarcinoma (CHOL), KIRP and PCPG (**Fig S1E; Table S1B**). Dysregulated IPAs are consistent among cancer types with similar histologic features (**Fig 1C**). For example, lung cancer (LUSC and LUAD), pan-kidney cancers (KIRC, KIRP and KICH), gynecologic and breast cancers (49) (uterine Corpus Endometrial Carcinoma [UCEC], CESC, and breast invasive carcinoma [BRCA]), show similar patterns in our hierarchical clustering analyses.

Although no IPAs have consistently higher or lower usage in tumors (vs. matched normal tissue) in all 22 cancer types, most (9,581 out of 11,975) show the same patterns in at least 2 cancer types (**Fig 1C**): 2,011 pan-cancer dysregulated IPAs with consistently higher or lower usage in at least 11 cancer types; 7,570 IPAs showing consistently higher or lower usage in 2 to 11 cancer types; and 2,394 cancer-specific IPAs that are dysregulated in only 1 cancer type. The pan-cancer dysregulated IPAs account for 52.1% of dysregulated IPAs in LIHC (the highest), but only 27.8% in CESC (the lowest); cancer-specific IPAs accounted for 9.2% in CESC and 1.6% in BLCA (highest and lowest, respectively) (**Fig S1F**).

### Pan-cancer IPA-derived early transcriptional termination

We defined one gene as a tumor-enriched or -depleted IPA gene based on the usage changes of IPAs within this gene (**Fig 2A & B**; **see Methods**). In the 690 tumors (compared to paired normal samples), these IPA-regulated genes vary from 132 to 2,249 (**Figs 2C & S2A**).

Many genes show consistent patterns of IPA regulation (**Fig S2B**). To study the consistency of IPA regulation, in each cancer type, one gene is defined as a cancer-type tumor-enriched IPA gene if more tumor samples (≥ 50 and at least 2) consistently exhibit this gene as a tumor-enriched IPA gene, or as a cancer-type tumor-depleted IPA gene if more samples (≥ 50 and at least 2) exhibit this gene as a tumor-depleted IPA gene (**Fig 2A; see Methods**). For example, RECK is a cancer-type tumor-enriched IPA-truncated gene found in 21 cancer samples, while ITFG2 is a cancer-type tumor-depleted IPA gene found in 19 cancer samples (**Fig 2B**). CESC has the most cancer-type tumor-enriched and -depleted genes and LIHC the fewest (**Table S2A**). Most cancer types show strikingly elevated number of cancer-type tumor-enriched than depleted IPA genes in the tumors, such as LUSC, BLCA, and LUAD; while KIRC, THCA, CHOL and KIRP have opposite trends (**Fig S2C; Table S2A**). In addition, unsupervised clustering analysis showed similarity of cancer-type tumor-enriched and -depleted genes among cancer types with similar histologic features (**Fig 2D**), such as lung cancer (LUSC and LUAD), pan-kidney cancers (KIRC, KIRP and KICH), pan-gyn organ sites (UCEC, CESC and BRCA), suggesting that IPA has a similar regulatory role in cancers of common tissue origin.

Our analyses yielded 4,491 cancer-type tumor-enriched or depleted IPA genes across 22 cancer types (**Fig 2D**). 771 genes (17.6%) consistently express the same transcripts in at least 11 cancer types, and 2,808 genes show consistent patterns in at least 2 cancer types. To further study the pan-cancer consistency, we identified 1,834 pan-cancer tumor-enriched IPA genes that occurred in at least half of the cancer types, which was about twice the number of pan-cancer tumor-depleted IPA genes (n=907) (**Fig 2D; Table S2B**). The pan-cancer genes account for 95% of IPA-regulated genes in BRCA (the highest), but 75% in PCPG (the lowest) (**Fig S2D**).

Next, we applied the bioinformatics tool CPAT (39) to predict coding probability of the IPA isoforms (**see Methods**). Of the 1,834 pan-cancer tumor-enriched IPA genes (**Fig S3A**), 791 IPA isoforms (43.6%) have coding probability scores greater than 0.364, meaning they have potential to produce protein-coding mRNAs and proteins (ee**Fig S3B**). The other 1043 IPAs (56.4%) (coding probability scores < 0.364) likely generate micropeptides or represent non-coding RNAs (IPA-ncRNA; **Fig S3C**), suggesting that IPA diversifies the transcriptomic landscape and results in potential loss of protein coding genes and their functional protein products in human cancers. Protein coding ability is not associated with polyadenylation signaling sequences, or IPA locations, but is associated with intron length (**Fig S3D**).

In addition, 453 IPA isoforms are in fact 5’ IPA isoforms (IPA located at first intron; **Fig S3D**). Of these, 212 may generate protein-coding genes with the length of open reading frames ranging between 126 bp and 5,394 bp (**Fig S3E**); others likely generate micropeptides or represent IPA-ncRNAs **(Fig S3F**).

### Pan-cancer tumor-enriched IPA genes are involved in cilium assembly and DNA repair

To better understand the biological consequences of IPA in cancer, we applied the enrichR tool (41) to characterize the pathways associated with the identified genes. Gene Ontology analyses showed that the 1,834 pan-cancer tumor-enriched IPA genes were enriched in the functional terms related to cilium assembly and DNA repair (**Fig 3A; Table S3A**). Evidence is accumulating that ciliary defects lead to multiple diseases (50) and ciliary deregulation plays crucial roles in cancer formation and progression (51-53); restoring the cilia can suppress proliferation in cancer cells (54,55).

**Figure 3:**
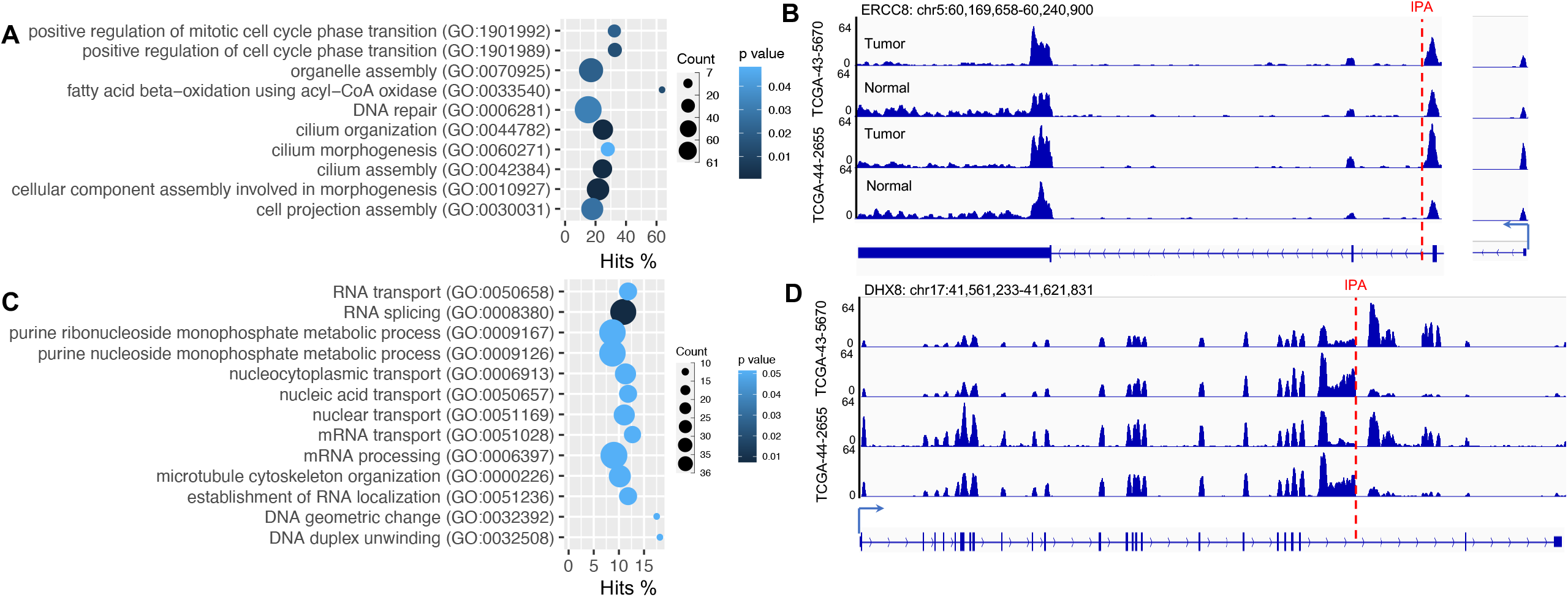
Functional enrichment of pan-cancer tumor-enriched and depleted IPA genes. (A,C) Gene Ontology enrichment of pan-cancer (A) tumor-enriched and (C) depleted IPA genes. (B,D) Examples of tumor-enriched (B: ERCC8) and depleted (D: DHX8) IPA DNA damage repair genes.

The DNA repair process operates through a number of mechanisms such as excision repair and homologous recombination repair to protect the human genome from damage and provide genome stability. DNA repair deficiency enables cancer cells to accumulate genomic alterations and contributes to their aggressive phenotype (56,57). Damage in one of the major DNA repair mechanisms, homologous recombination, by IPA has been reported previously through the depletion of CDK12 (27,28). ERCC8 (**Fig 3B**) is the top gene that is truncated in tumors by aberrant IPAs in 17 cancer types, followed by FANCN. The former participates in excision repair and the latter plays roles in double-strand break repair.

The 904 pan-cancer tumor-depleted IPA genes that preferentially produce full-length mRNAs in tumors are enriched in mRNA processing tasks such as splicing and 3’-end processing (**Fig 3C; Table S3B**). For example, the IPAs of DHX8, encoding an RNA-binding protein involved in splicing, are inhibited in 13 cancer types (**Fig 3D**), and the IPAs of hnRNPL, encoding a protein regulating both mRNA splicing and IPA processing (58,59), are inhibited in 11 cancer types. These data indicate that IPA truncation of these genes is inhibited in tumors, which, in turn, promotes the post-transcriptional regulation of transcriptomics program in human cancers. Genes associated with cancer hallmark signatures, such as epithelial-mesenchymal transition and E2F targets, are also overrepresented in this group of genes (**Fig S4**), suggesting that IPA inhibition in these genes have an oncogenic effect.

### Correlations between truncating mutations, IPA, and tumor mutation burden

We showed that the aberrant IPAs in tumors caused early termination of transcription of many genes, some of which generated truncated proteins that lack essential domains or IPA-ncRNAs. This phenocopied the effects of truncating mutations at DNA level. Overall, 563 (out of 690 in 22 cancer types) samples carry truncating mutations in 5,344 genes. By contrast, truncation by aberrant IPAs occurs in all samples and 6,055 genes. Only 377 genes in 148 samples from 17 cancer types carry both truncating mutation and aberrant IPAs (**Table S4A**), suggesting that these two actions independently regulate transcriptome diversity. In addition, in all individual samples, more genes are affected by IPAs than truncating mutations (**Fig 4A; Table S4B**).

**Figure 4:**
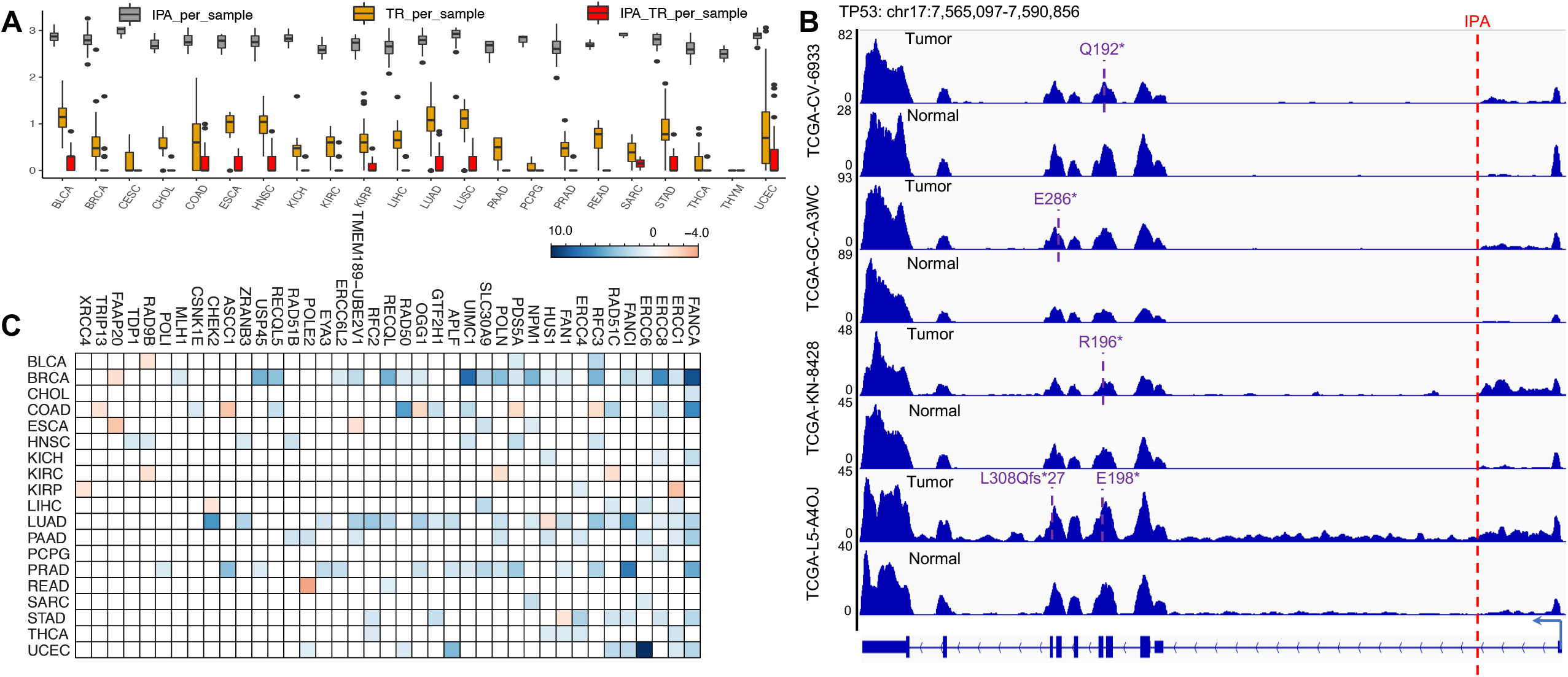
IPA and truncating mutations independently diverse cancer transcriptomics. (A) Numbers of genes truncated by IPA, truncating mutations, or both in each cancer type. (B) Samples having both TP53 truncating mutations and IPA isoforms in tumors compared to matched normal samples. Pan-cancer tumor-enriched IPA DNA repair genes associated with tumor mutation burden. Colors indicate the log10(p value) with positive (blue) or negative (orange) correlations.

We then examined DNA damage repair genes in particular, since the loss of DNA repair has been identified in many cancer types due to somatic mutations (60). 132 DNA damage repair genes exhibit aberrant IPAs in all 690 samples, and 77 genes have truncating mutations in 94 samples; 11 genes have both in at least one sample (**Table S4B**). RAD51B is the top gene regulated only by aberrant IPAs in 425 samples (out of 690; 61.5%). TP53 is the top gene carrying truncating mutations (32 samples; 4.6%), existing in BRCA (n=2), colon adenocarcinoma (COAD; n=2), ESCA (n=1), head and neck squamous cell carcinoma (HNSC; n=10), KICH (n=1), LUAD (n=6), LUSC (n=3), rectum adenocarcinoma (READ; n=2) and stomach adenocarcinoma (STAD; n=2), or both (4 samples from KICH, ESCA, HNSC and BLCA; 0.58%) (**Fig 4B; Table S4C**). In contrast, MEN1 is the most frequently truncating mutation-only gene, which occurs in only 3 samples (from HNSC, LUAD and STAD).

Because genes targeted by truncating mutations are often tumor suppressor genes (42), we also investigated whether such genes are overrepresented among those regulated by IPAs. Among 301 tumor suppressor genes (42), 62 do not contain detectable IPAs or truncating mutations. Among the other 239 genes, 151 genes express IPA-truncated IPA isoforms in at least one of the 690 samples, and 191 genes contain truncating mutations. FUBP1 is the top 2^nd^ gene regulated only by IPA in 419 samples (out of 690; 60.7%), following TP53 that is defined as both DNA damage repair and tumor suppressor genes; APC is the top gene regulated by truncating mutations (21 samples; 3.0%). Only 27 genes have co-occurring IPAs and truncating mutations, including KMT2C (5 samples), TP53 (4 samples), CASP8 (3 samples), PTEN (3 samples), NSD1 (2 samples) and SETD2 (2 samples) (**Fig S5A; Table S4D**). KMT2C (aka MLL3) is a chromatin remodeling gene that mediate H3K4 monomethylation (H3K4me1). KMT2C mutations contribute to tumorigenesis and are associated with poor survival rates (61,62). In particular, the complete loss of KMT2C due to truncating mutations is correlated with significantly shorter progression-free survival in patients with breast cancer patients who undergo estrogen therapy (63) and in patients with metastatic prostate cancer (64). The IPA-associated loss of KMT2C may have the functions that need further validation.

Next, we tested whether IPA isoform expression is associated with tumor mutation burden, since impaired DNA damage repair often leads to abundant mutations (65). In this analysis, all 8,039 samples of the 22 cancer types are included (**see Methods**). Most pan-cancer tumor-enriched IPA genes (n=1,641) have significant associations with tumor mutation burden in at least one cancer type (**Table S4E**). Across the 22 cancer types, BRCA has the most IPAs associated with tumor mutation burden (n=903), followed by LUAD (n=650) and prostate adenocarcinoma (PRAD; n=447); PCPG (n=35) and CHOL (n=30) have the fewest IPAs associated with tumor mutation burden. Also, most cancer types show strikingly more positively correlated IPA isoforms, such as LUAD and pancreatic adenocarcinoma (PAAD) (**Fig S5B**). Specifically, 39 (out of 61) pan-cancer tumor-enriched IPA isoforms of DNA damage repair genes are associated with tumor mutation burden, most of which are positively associated across cancer types (**Fig 4C**). For example, FNACA is a member of the Fanconi anemia complementation group that is involved in both DNA interstrand crosslink repair and double-strand break repair. The expression of its IPA isoform is positively correlated with tumor mutation burden in 9 cancer types (**Fig S5C**). Therefore, impairment of DNA repair genes by IPA events may be a major cause of mutation accumulation in human tumors.

### Upstream regulatory factors for IPAs and effects of IPA-ncRNAs on signaling pathways

To investigate the potential regulatory factors for IPA in human cancer, given the interplay between alternative splicing and intronic polyadenylation, we examined 42 polyadenylation factors and 262 splicing factors. 8,039 samples of the 22 cancer types are included in analysis. Positive and negative regulators are defined as whether one regulator is positively or negatively correlated with more IPAs in a given cancer type (**see Methods**). Taking HNSC as an example, we identified 190 putative master regulators, including 108 positive and 82 negative regulators.

Applying this computational approach, we identified 282 putative master regulators across cancer types (**Table S5**), among which 95 and 59 factors exhibited strong positive and negative correlation with multiple IPAs in 11∼22 cancer types, and 30 and 11 factors showed correlations in only 1 cancer type. CPSF3 is the top polyadenylation factor that broadly promotes IPA activities in almost all cancer lineages, ranging from 24% in PCPG to 94% in LUSC (**Figs S6A & S6B**). The splicing factors POLR2D, EIF4A3, and HSPA8 are the top master regulators positively correlated with more IPAs across all cancers (**Fig S6C**).

Among the 282 master regulators, we identified 18 as pan-cancer tumor-enriched IPA genes and 29 as pan-cancer tumor-depleted IPA genes, respectively (**Fig 5**). PCF11 is an tumor-enriched IPA gene whose transcription is terminated by IPA in 7 cancer types (**Fig S6D**), where its gene expression is inversively corrleated with IPA isoform expression (**Fig S6E**). hnRNPL is an tumor-depleted IPA gene whose full-length transcript is consistently expressed in 11 cancer types, and its gene expression is positively correlated with IPA isoform expression. These data are consistent with the previously reported roles of PCF11 (66) and hnRNPL (58) in the IPA process.

**Figure 5:**
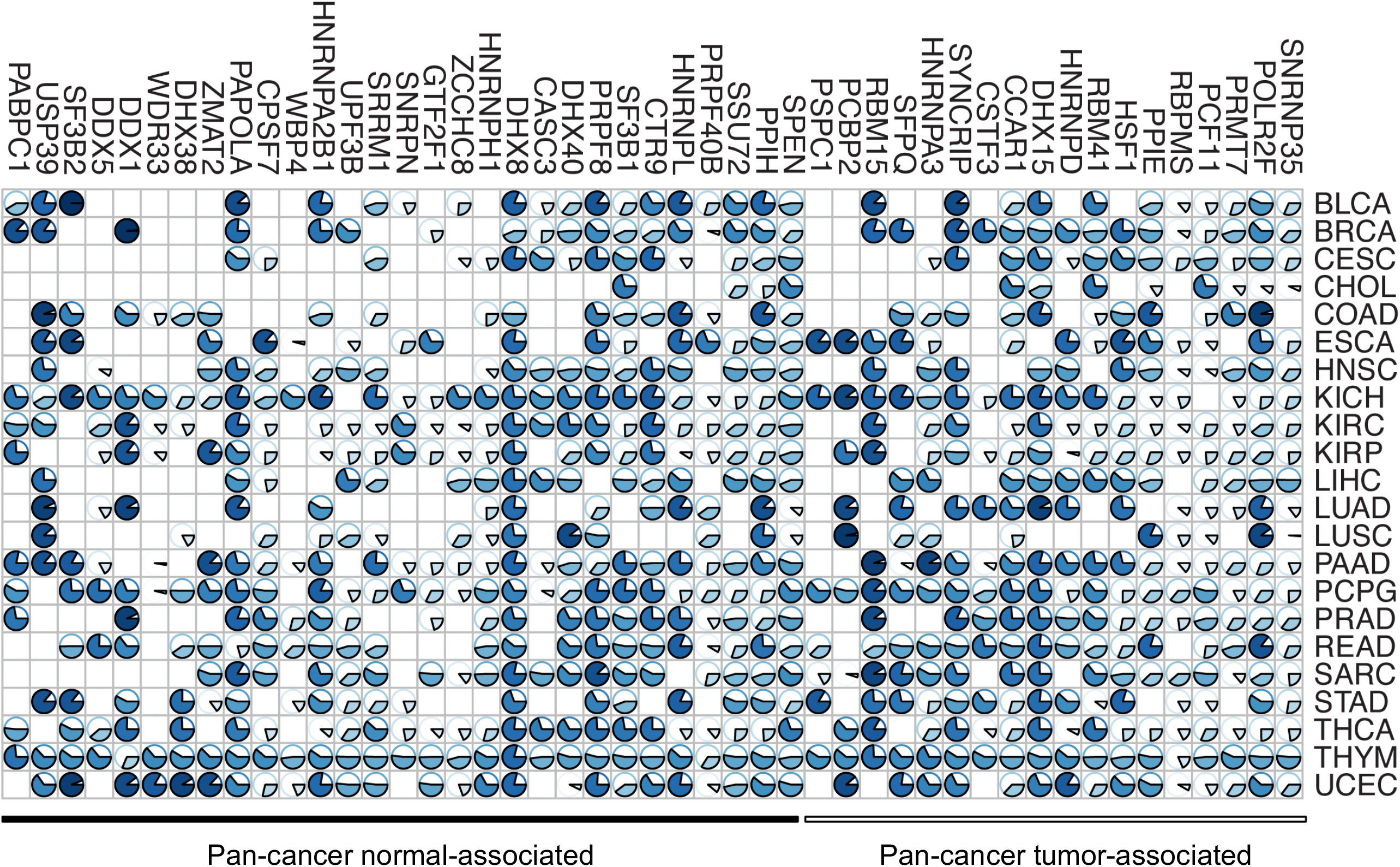
Putative regulators of IPA activities in human cancers. Pie charts show percentage of IPA isoforms that are positively (blue) or negatively (white) correlated with regulators (that were defined as pan-cancer tumor-enriched or depleted) in each cancer type.

ncRNAs play critical roles in cellular processes related to tumorigenesis by interacting with mRNAs (67-71), and may help inform optimal drug design (72,73). We examined interactions between 1,043 IPA-ncRNA isoforms determined previously (coding probability score < 0.364) and their associated genes. We focused on the associations of these IPA-ncRNAs and 335 genes involved in 10 cancer signaling pathways (p53, PI3K, Myc, RTK/RAS, cell cycle, Wnt, TGF beta, Nrf2, Notch, and Hippo) (47), and built an IPA ncRNA-gene regulatory network across cancer types based on co-expression between individual IPA and their putative target genes (**see Methods**). We found that 735 IPA-ncRNAs were computationally correlated with 327 target genes in the 22 cancer types. The most strongly associated pathway is RTK/RAS, especially in THYM and PAAD (**Fig S7A**), and the top associated gene is RAC1 in the RTK/RAS pathway in PAAD, which is associated with expression of 98 IPA-ncRNAs (**Fig S7B, Table S6**), such as the IPA-ncRNA from TMEM8B (**Fig S7C**).

### Clinical relevance of IPA isoforms

We hypothesized that individual variations in IPA isoform expression might provide a meaningful predictor for patient clinical outcomes. Within the TCGA cohort, we categorized patients into two groups (≥ or < 90%; **see Methods**) based on levels of IPA expression or gene expression (sum of all isoforms e.g. those from alternative polyadenylation and splicing) of each gene. Among the 1,834 pan-cancer tumor-enriched IPA genes, many show significant associations with overall survival for either IPA isoform or total gene expression (the sum of all possible isoforms), but not both (**Fig 6A; Table S7A & S7B**). The largest gene set (n = 225) for which gene expression and IPA isoform expression predicted patient survival was found in KIRC samples, whereas only 1 gene was found in the THYM. Intrudingly, we found 225 genes whose expression and IPA isoforms predicted opposite survival outcomes in at least one cancer type (**Table S7B**). For example, KDM4C has been reported as an oncogene and a negative biomarker for patients with multiple cancer types (74-76), and is associated with cancer metastasis (77) and immunosuppressive tumor microenvironment (78). However, its gene expression is associated with better survival outcomes in patients with LUAD – unlike expression of its IPA isoform, which is associated with poor survival of those patients (**Figs 6B & S8A**). This negative correlation is also observed in patients with LUSC, KICH and BRCA, where the gene expression does not show significant association (**Table S7B**). Another example is DNAJB4 (HLJ1); although reported as a tumor suppressor gene (79,80), we found negative associations between its gene expression level and patient survival in multiple cancer types (**Fig 6B**; **Table S7B**). However, its IPA isoform expression is a biomarker of better survival (**Figs 6B** & **S8B**). These observations suggest consideration of gene isoforms in cancer biomarker studies.

**Figure 6:**
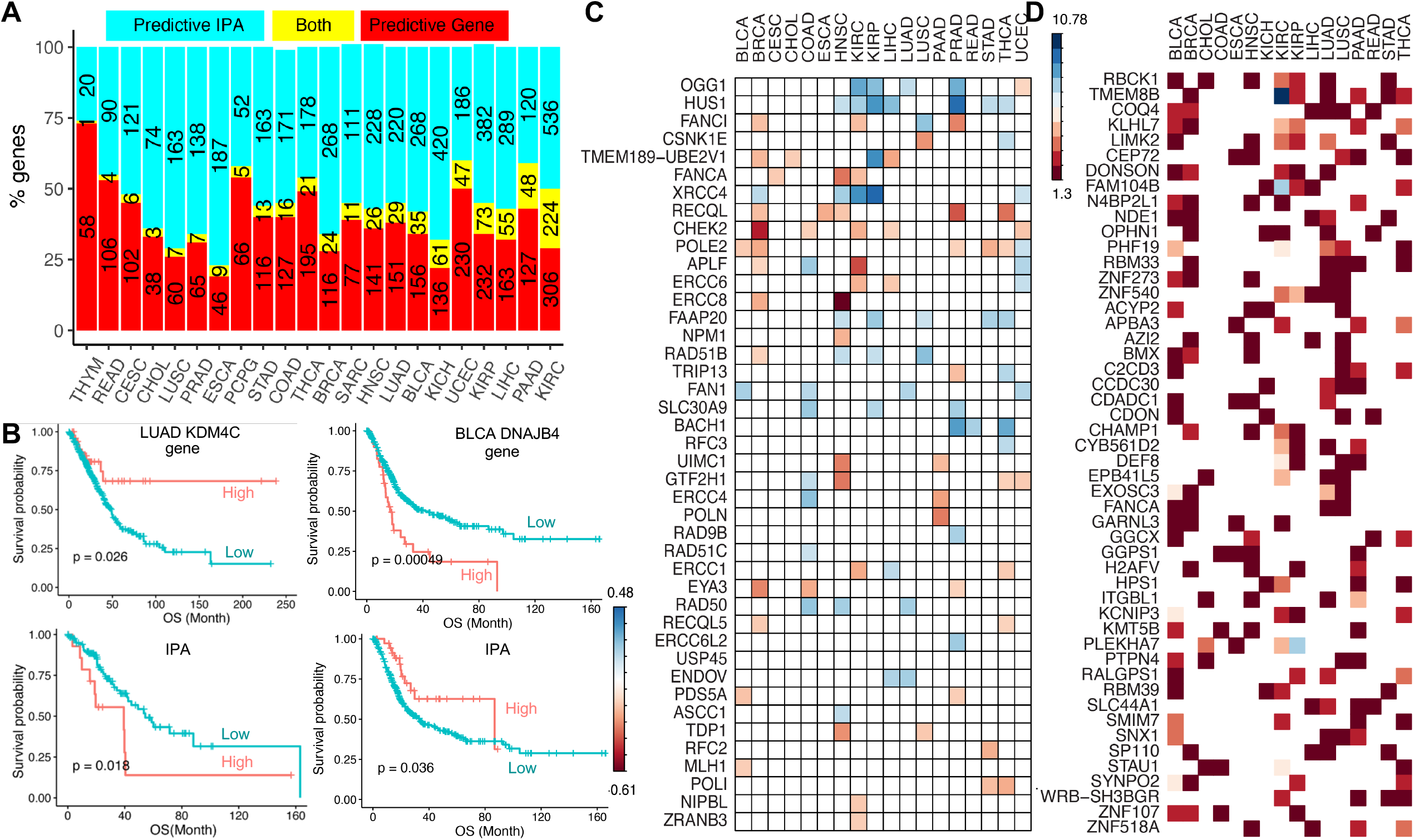
Associations between IPA isoforms and clinical characteristics. (A) Numbers of genes whose expression or IPA usage predicted patient survival. (B) IPA isoform and gene expression of KDM4C and DNAJB4 had opposite correlations with overall survival in the TCGA cohort. (C) Pan-cancer tumor-enriched IPA DNA repair genes and associations with CD8 cell fractions. Colors indicated the log10(p value) with positive (blue) or negative (orange) correlations. (D) The top 50 genes showing significant associations with cancer stages in the TCGA cohort. Colors indicated the log10(p value)

Next, we tested whether IPAs were associated with patient immune cell profiles. The immune cell fractions of patient tumors in the TCGA cohort are estimated using CIBERSORT (46) (**see Methods**). Among the 61 pan-cancer tumor-enriched IPA isoforms in DNA damage repair genes, 39 are consistently associated with tumor immune cell profiles across cancer types (**Table S7C**). The expression of HUS1 IPA isoform is positively correlated with CD8 T cell populations in 7 cancer types (**Fig 6C**) and with natural killer cell populations in 6 cancer types (**Fig S8C**), indicating that the truncation of HUS1 transcription (and resultant decrease in levels of HUS1 functional proteins) promotes an anti-tumor tumor microenvironment. This observation is consistent with previous findings that depletion of HUS1 by miRNA inhibits *de novo* lung tumorigenesis (81). In a pan-cancer study, variant or low expression levels of FANCI are associated with poor patient survival (82). This is consistent with our pan-cancer observations that expression of FANCI IPA isoform expression is negatively correlated with CD8T cells (**Fig 6C**), activated CD4T cells, and natural killer cells (**Fig S8C**). XRCC4 is among the canonical-nonhomologous end-joining (C-NHEJ) DNA repair components. C-NHEJ deficiency is associated with severe combined immunodeficiency and accelerated tumor development (83). Our analysis also revealed that high levels of XRCC4 IPA isoform expression were consistently associated with regulatory T cell populations in 9 cancer types (**Table S7C**).

We then selected cancer-related demographic and clinical features, including stages (available for 16 cancer types), race (available for 22 cancer types), sex (available for 15 cancer types; cancers specific to men or women were excluded), and subtypes (available for 9 cancer types) (**Table S7D**), and detected associations between IPAs and these factors. Overall, we identified 1,371 clinically relevant IPA isoforms that accounted for 74.8% (1,371/1,834) of pan-cancer tumor-enriched IPA genes (**Table S7E**), including 820 associated with cancer stage, 810 associated with patient race, 557 differently expressed in male and female patients, and 1,100 correlated with cancer subtypes. Most IPA isoforms show cancer-specific patterns regarding individual clinical features. Among those, 461 (56.2% of 820) IPA-cancer stage associations occur in only 1 cancer type; in contrast, no gene show associations in more than 8 (50% of 16) cancer types. The top genes are RBCK1 and TMEM8B, which are associated with tumor stages in 7 cancer types (**Fig 6D**). 411 (50.7% of 810) IPAs are associated with patient races in only 1 cancer type, while no gene showed associations in more than 11 (50% of 15) cancer types. The most highly race-associated genes are BOD1L1 and ELP6 in 9 cancer types (**Fig S8D**). 519 (93.2% of 557) IPAs are associated with patient sex in only 1 cancer type, while only one gene (KDM5C) has associations with more than 8 (50% of 22) cancer types (**Fig S8E**). 433 (39.4% of 1,100) IPA are associated with patient subtypes in only 1 cancer type, and 89 genes show associations in more than 5 (50% of 9) cancer types. Two genes, PDK3 and ZFAND4, have significant associations in 8 cancer types (**Fig S8F**).

In addition, most IPAs show feature-specific patterns within individual cancer types (**Fig S8G**). Five cancer types (COAD, ESCA, HNSC, READ and STAD) have all 4 features available, but only 4 genes in 3 cancer types, including BOD1L1 and RGS5 in COAD, GGCX in HNSC and SHANK3 in STAD, are associated with all 4 features, in contrast to 220 (73.3% of 300) in COAD, 165 (80.1% of 204) in ESCA, 203 (65.5% of 310) in HNSC 126 genes (86.9% of 145) in READ and 213 (72.7% of 293) in STAD associated with only 1 features. 11 cancer types (BLCA, BRCA, CHOL, KICH, KIRC, KIRP, LIHC, LUAD, LUSC, PAAD and THCA) have 3 features available, but only 66 genes (15.2% of 433) in BRCA, 8 (3.2% of 252) in KIRC, 3 (1.5% of 204) in KIRP, 6 (2.5% of 236) in LUAD, 5 (1.6% of 307) in LUSC and 1 (0.5% of 191) in THCA are associated with all 3 features. In contrast, most genes are associated with only 1 factor with number varying between 65 (95.6% of 68 in LIHC) and 244 (79.5% of 307 in LUSC). We observed similar patterns in the three cancer types with only 2 features available, 40 vs. 343 genes in CESC, 17 vs. 384 in sarcoma (SARC) and 131 vs. 312 in UCEC showing associations with 2 and 1 features, respectively.

## DISCUSSION

Advances in high-throughput transcriptomic analysis have revealed abundant RNA post-transcriptional processing mechanisms, including IPA (27-30). Interrogating large-scale RNAseq data sets from the TCGA cohort, we provided a systemic view of the IPA landscape in human cancer. We demonstrated a strong pan-cancer pattern of aberrent IPAs and IPA-regulated genes. These pan-cancer turmor-enriched IPA genes are enriched in functional pathways such as DNA repair, suggesting that dysregulated IPAs are involved in accumulation of somatic mutations in human tumors. This concept is supported by our observations that many IPAs are positively correlated with patient tumor mutation burden. In addition, expression of turmor-enriched IPA isoforms is associated with patients’ clinical characteristics such as survival, immune profiles, stage, subtype, sex, and race, suggesting the importance of IPA isoform independent of gene expression (the sum of all isoforms).

Our analysis identified 22,260 detectable IPA sites, most of which showed a consistent expression pattern across 33 cancer types. Pan-cancer IPAs (**Fig 1C**) have higher usage levels (Wilcoxon test *p* < 2.2 × 10^−16^) than others, which is reminiscent of features of protein-coding genes and eRNAs -- the housekeeping genes and pan-cancer eRNAs are generally expressed at high levels compared with tissue-specific genes (84) and eRNAs (85), respectively.

Tumor cells express transcripts with systematically shorter 3′ UTRs compared to normal cells (14) by selecting the proximal polyadenylation sites in 3’UTR; many oncogenes are upregulated through this mechanism by escaping regulatory miRNA-mediated repression (10,13,86). We observed numerous tumor-enriched and depleted IPA genes in each tumor (compared to the paired normal tissues). Tumors also tended to express IPA-truncated isoforms; more samples had a global IPA-related early transcriptional termination (n = 460 and 230, respectively). Unlike 3’UTR-APA, which mainly functions in oncogenes, we found that pan-cancer tumor-enriched IPA genes were largely associated with DNA repair pathways, and tumor-depleted IPA genes were associated with cancer hallmark signatures such as EMT and E2F targets. Therefore, APA processing, including both 3’UTR-APA and IPA, may participate in tumor development and progression, suggesting its potential as a treatment strategy.

Both truncating mutations and aberrant IPAs in genes cause early transcriptional termination of protein-coding gene transcription and generate non-coding RNAs or dysfunctional mRNAs. We observed that IPA regulation exclusively occurred with truncating mutations, especially in the DNA repair and tumor suppressor genes; loss of those genes has been linked to somatic truncating mutations in human cancer genes (42,60). Therefore, IPA, in addition to somatic mutations, diversifies the cancer transcriptomic landscape and hampers the tumor suppressive mechanism in human cancers.

The importance of long noncoding RNAs (lncRNAs) has just recently been appreciated and is still debated. LncRNAs may play roles in different mechanisms of gene regulation. Increasingly, lncRNAs are considered important in cancer initiation and progression, patient drug resistance and treatment outcomes, and potential therapeutic targets (87). Our analysis revealed that IPA truncation may result in many IPA-ncRNAs. These can regulate gene expression and alter cancer signaling pathway activities. Further, we identified an appreciable number of IPA isoforms with potential clinical relevance that could be potential biomarkers. Gene expression (the sum of all possible isoforms, e.g. those from alternative splicing and polyadenylations) has been employed for the biomarker identification, but this approach may not be accurate due to the possiblely different functions of individual isoforms. We identified a set of genes whose IPA isoform expression and total gene expression were conversely associated with patient survival, e.g. KDM4C and DNAJB4 (**Fig 6B**). This observation suggests that isoform expression is more accurate prognostic marker than gene expression.

DNA damage repair (DDR) and the immune system are tightly connected. DDR deficiency caused by somatic mutations regulates the innate and anti-tumor adaptive immune system through multiple pathways, e.g. the cyclic GMP–AMP synthase (cGAS)-stimulator of interferon genes (STING) pathway (88). Mutations in DNA repair genes and elevated tumor mutation burden are associated with cancer immunotherapeutic response and cancer prognosis (89-91). IPA truncation of these genes also enables DDR deficiency; as a result, we found that DDR IPA isoform expression was strongly associated with immune cell profiles in patient tumors. The IPA-induced truncation of transcription may produce neoantigens, similar to mRNA splicing or truncating mutations. Neoantigens can stimulate an immune system response against cancer cells (92,93) and is a promising approach to tumor immunotherapy (94,95).

Our study does have limitations. For example, we acknowledge the limitations of using bulk RNA-seq data for studying the tumor microenvironment and anti-tumor immunity. Detailed functional studies of IPA in cancer immunity will require more refined techniques, such as single-cell RNA sequencing. Although our analysis used the largest human cancer cohort, many molecular measurements, such as miRNA expression and DNA methylation, were not included.

In summary, by using large-scale RNAseq and clinical datasets from the TCGA cohort, we provide a comprehensive landscape of IPA events in human tumors and their clinical relevence. Our analysis can be a useful resource for further investigation regarding regulating mechanisms and functions of IPA.

## ACKNOWLEDGEMENT

We would like to acknowledge the editorial assistance of Karen Klein, MA, ELS, MWC, from the Clarus Editorial Services.

## FIGURE LEGENDS

**Figure S1.**
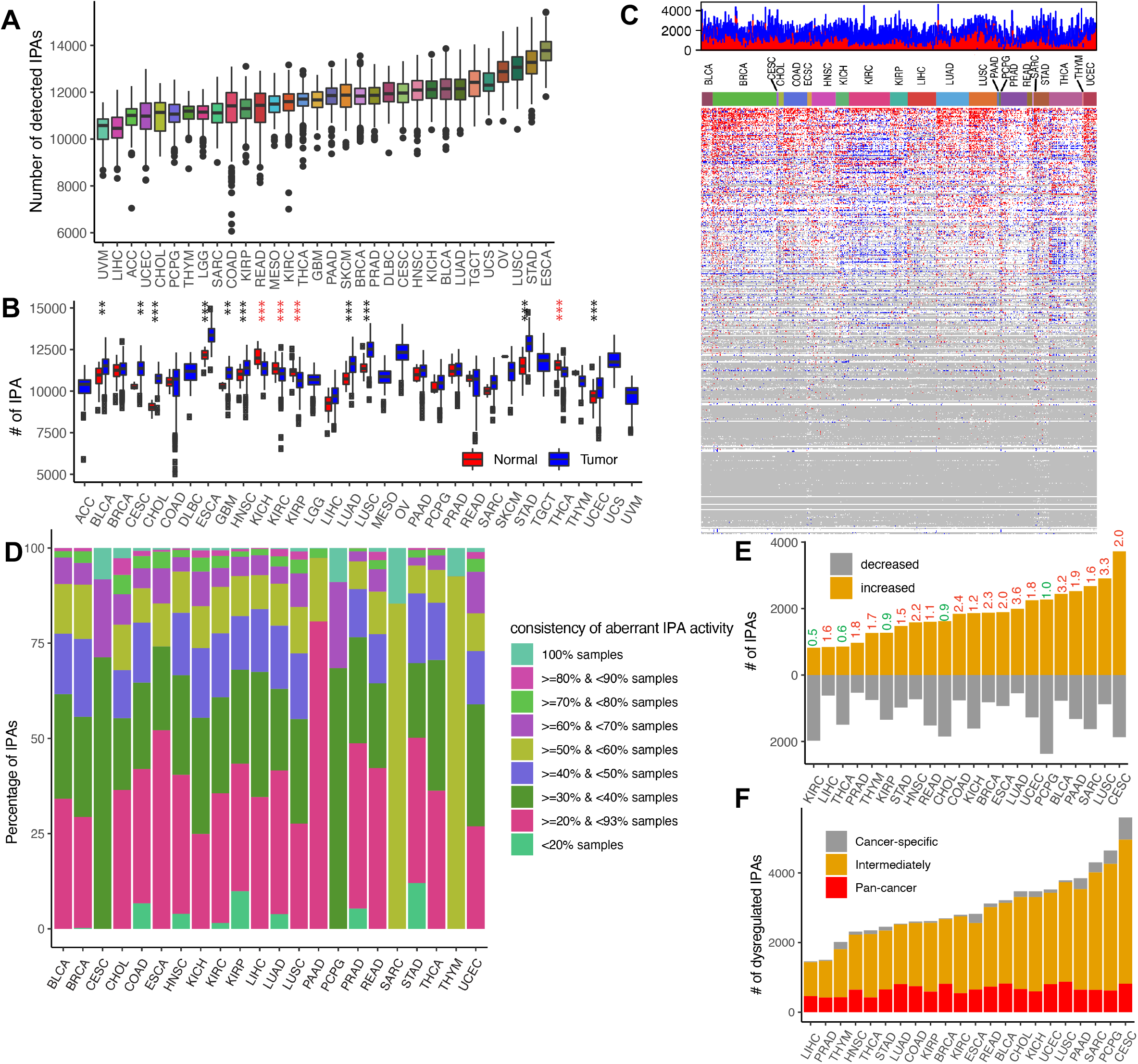
(A) Numbers of detectable IPAs in each cancer type; related to Fig 1C. (B) IPA numbers in tumor versus normal samples in each cancer type. (C) The central heatmap shows IPAs (row) with higher (red) or lower (blue) usage in each tumor (columns) compared with matched normal tissue across tumor types. The upper histogram shows the number of IPA events per tumor. (D) Consistency of IPA regulation in individual cancer types. (E) Numbers of IPAs with higher or lower usage (tumor vs. matched normal) in each cancer type. Ratio values (number of IPAs with higher/lower usage) were shown for each cancer type. (F) Percentages of dysregulated IPAs in the three groups (tumor vs. matched normal).

**Figure S2.**
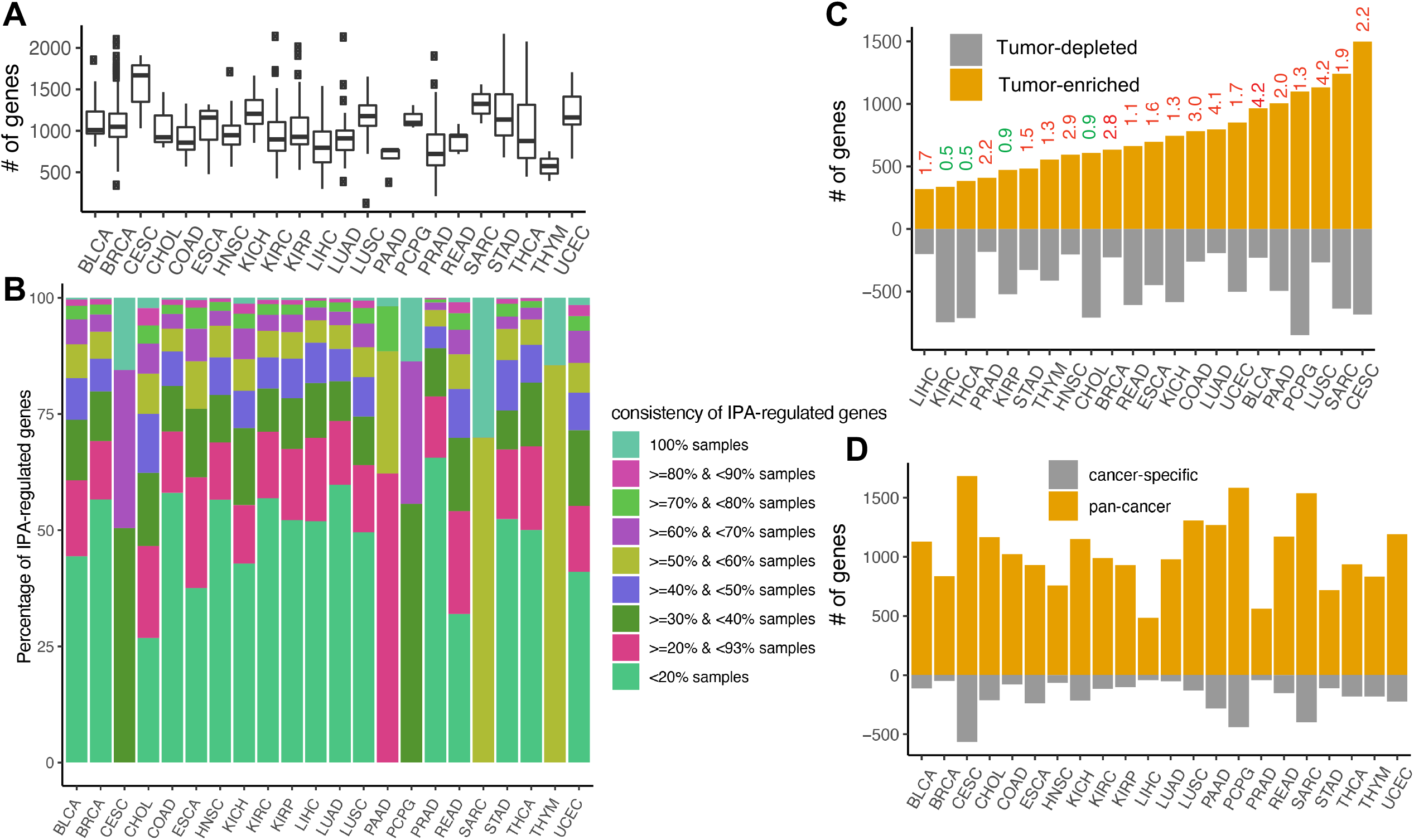
(A) Genes regulated by dysregulated IPAs in each cancer type. (B) Consistency of IPA-regulated genes in individual cancer types. (C) Genes defined as tumor-enriched or depleted IPA genes in each cancer type. The ratio values (# tumor-enriched/# tumor-depleted genes) was shown for each cancer type. (D) Number of pan-cancer and cancer-specific IPA-regulated genes across cancer types.

**Figure S3:**
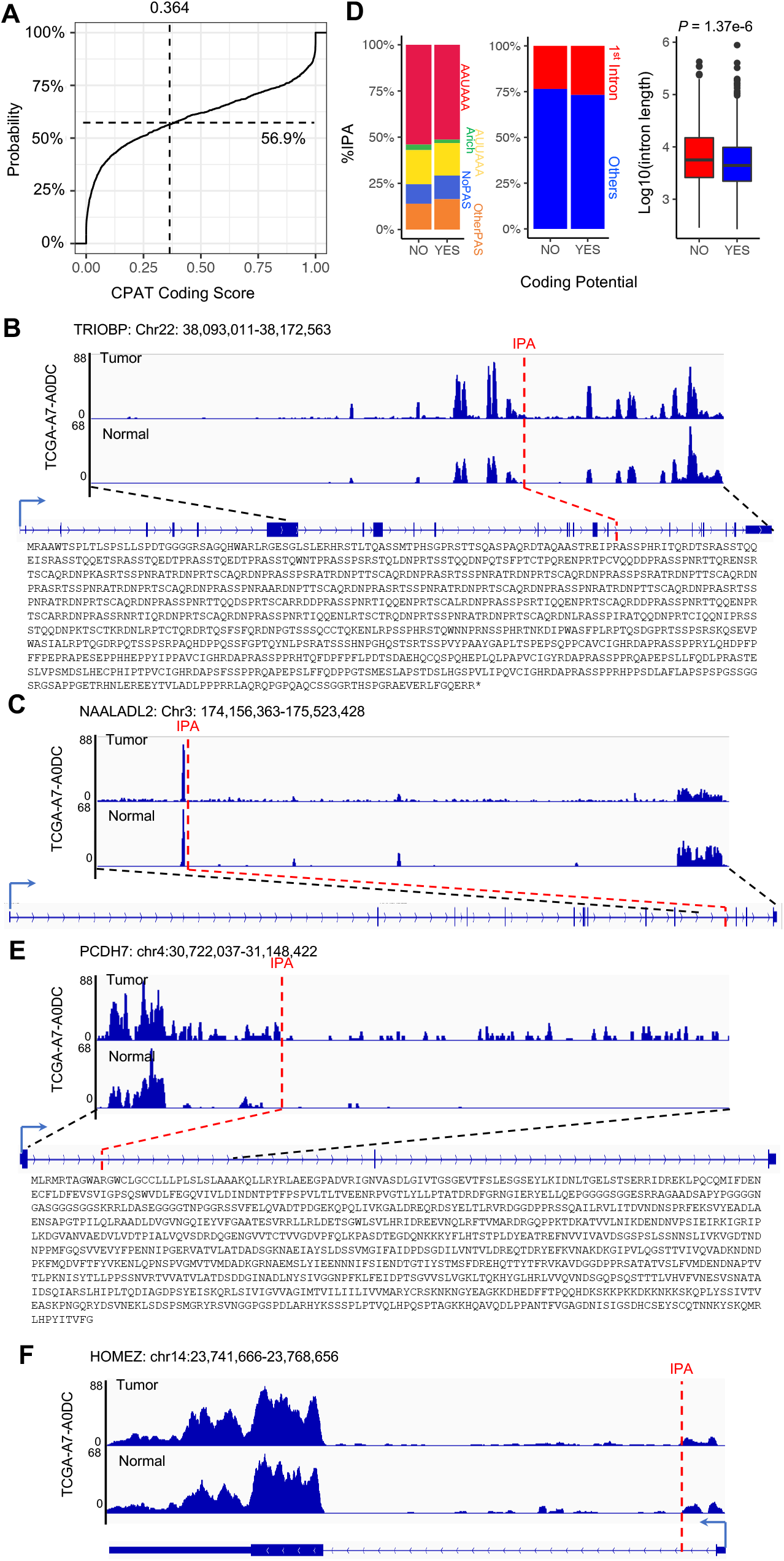
Products of tumor-enriched IPA genes. (A) CPAT coding scores of 1,834 pan-cancer tumor-enriched IPA gene isoforms. (B,C) Examples of IPA isoforms that produce (B) coding and (C) non-coding RNAs. The protein sequence was predicted using EMBOSS Transeq. (D) Correlations of polyadenylation signaling sequences, intron locations, and intron lengths with IPA isoform protein-coding abilities. (E,F) Examples of 5’IPA isoforms that produce (E) coding and (F) non-coding RNAs.

**Figure S4:**
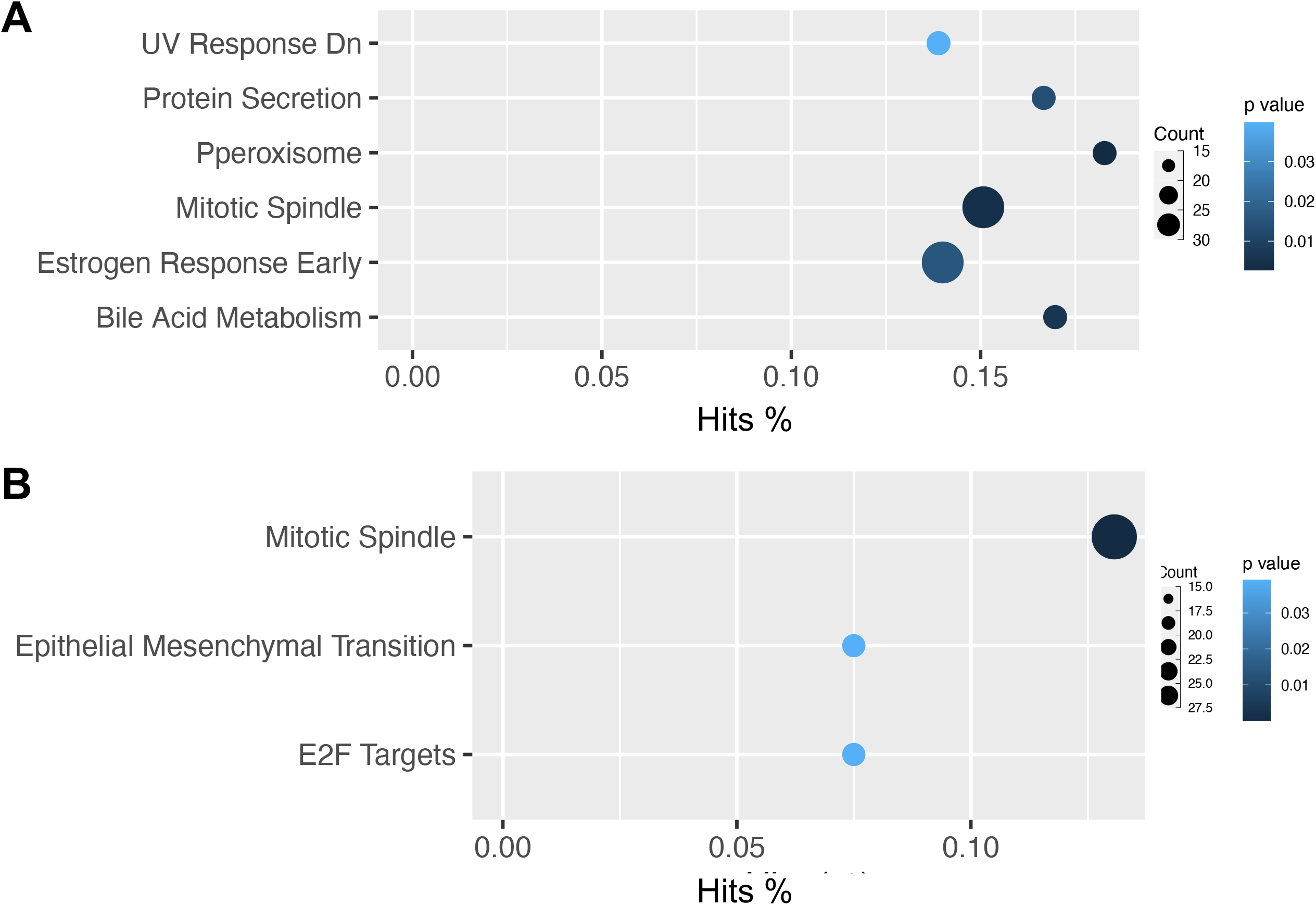
Hallmark signature enrichment of pan-cancer (A) tumor-enriched and (B) depleted IPA genes.

**Figure S5:**
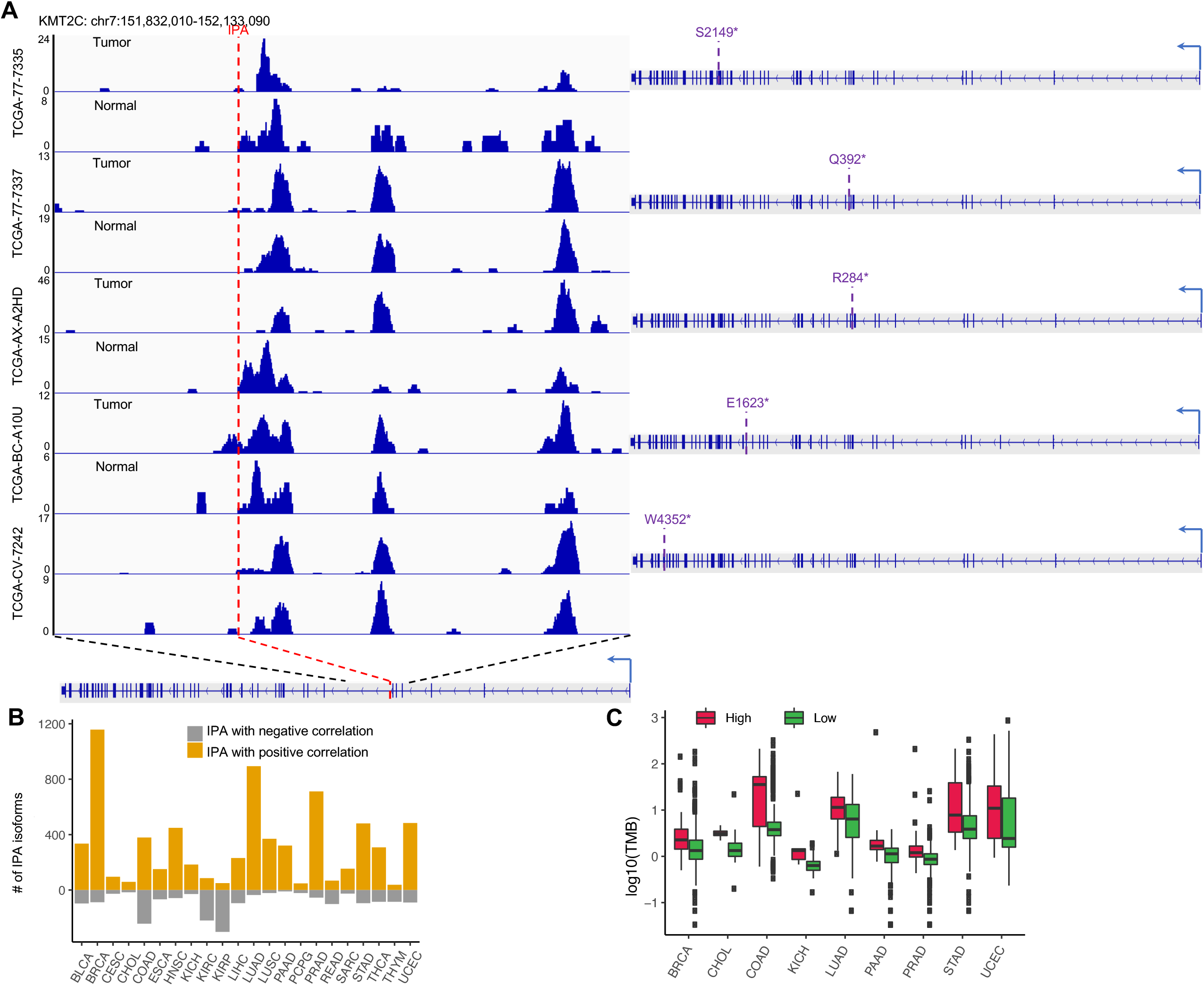
(A) Samples carrying both KMT2C truncating mutations and IPA isoforms in tumors compared to matched normal. (B) Numbers of IPA isoforms positively or negatively associated with tumor mutation burden in each cancer type. (C) IPA isoform of FANCA was positively associated with tumor mutation burden in 9 cancer types (Wilcoxon test *P* < 0.05).

**Figure S6:**
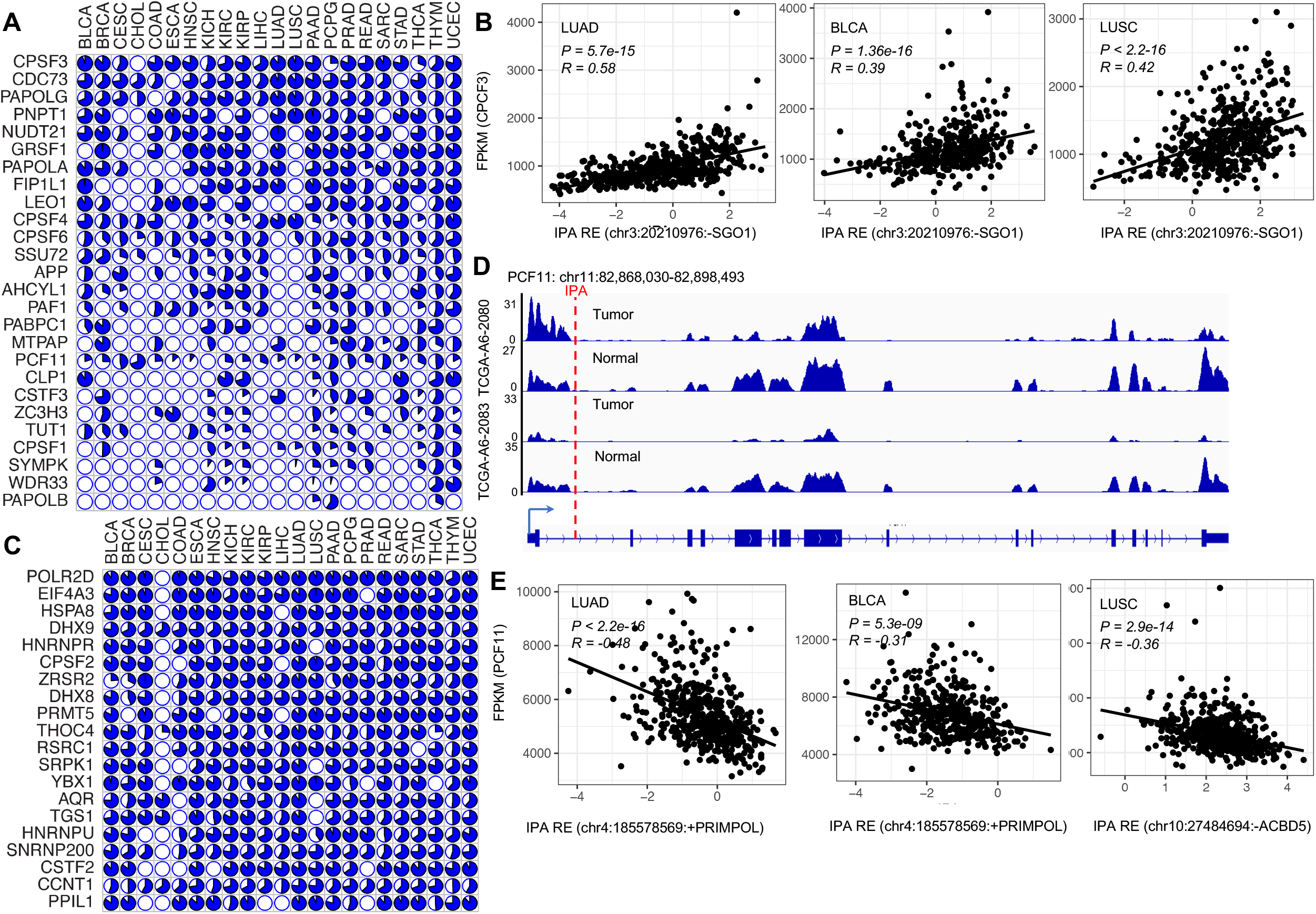
(A,C) Correlations between IPA isoform expression and (A) polyadenylation and (C) splicing factors. Pie charts show percentage of IPA isoforms positively (blue) or negatively (white) correlated with regulators in each cancer type. (B) Correlations between CPCF3 and IPA isoforms in different cancer types. (D) IPA isoforms of PCF11 enriched in tumors. (E) Correlations between PCF11 and IPA isoforms in different cancer types.

**Figure S7:**
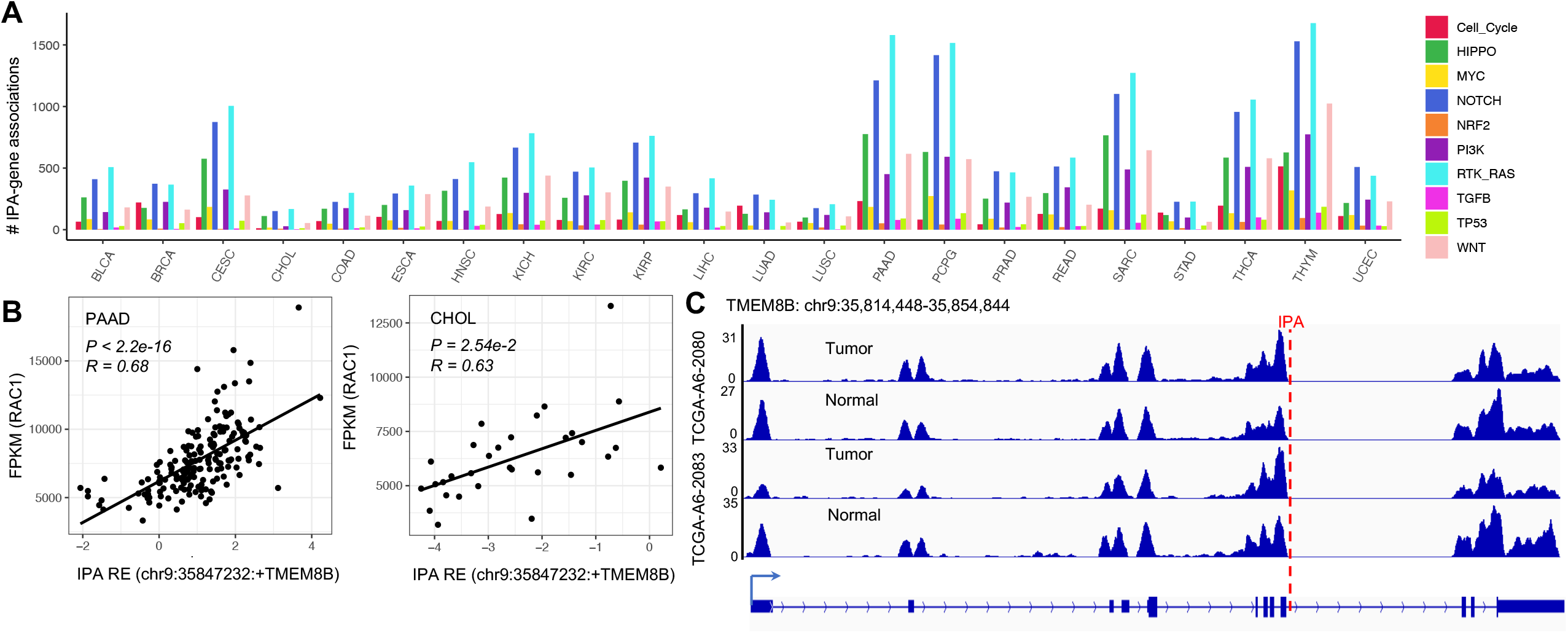
Putatively regulatory network of IPA-ncRNAs in signaling pathways. (A) IPA-gene associated pairs across pathways and cancer types. (B) Correlations between RAC1 expression and TMEM8B IPA isoforms. (C) Example of TMEM8B IPA isoforms.

**Figure S8:**
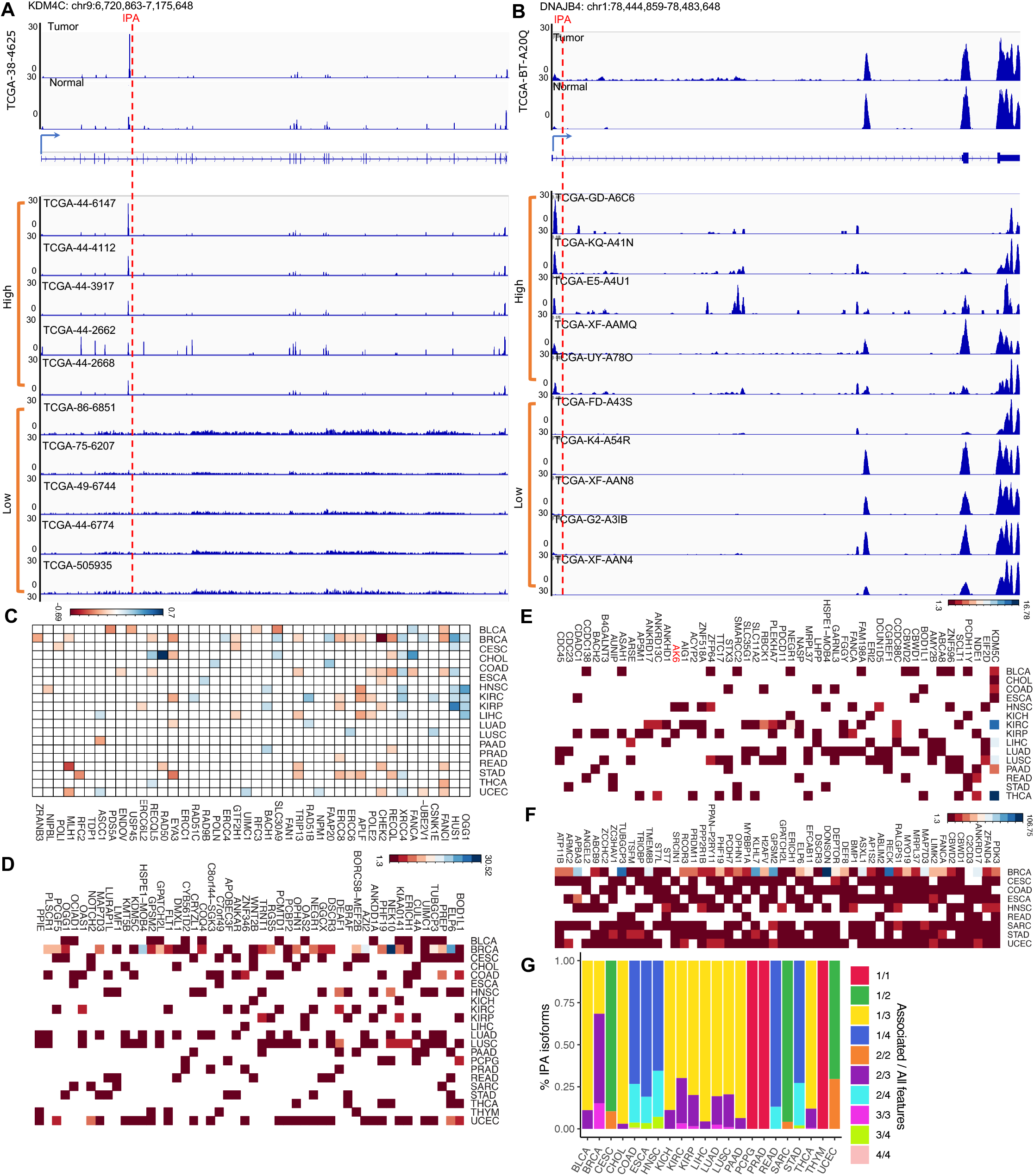
(A,B) RNAseq examples show (A) KMT4C and (B) IPA isoform is increased expressed in tumors, and samples with high and low KDM4C IPA isoform expression. (C) Pan-cancer tumor-associated IPA-truncated DNA repair genes associated with natural killer cell fractions. Colors indicated the log10(p value) with positive (blue) or negative (orange) correlations. (D-F) The top 50 genes show significant associations with clinical features, including (D) race, (E) sex, and (F) tumor subtypes. Colors indicate the correlation p values (-log10 transferred). (G) Percentages of genes associated with associated features / all available features.

## SUPPLEMENTAL TABLE LEGENDS

**Table S1:** (A) Cancer abbreviation in TCGA cohort.

(B) The number of tumors with matched normals, and the IPAs with significantly different usage in tumor vs. normal.

**Table S2:** (A) The number of tumors with matched normals, and the tumor enriched or depleted IPA-truncated genes.

(B) List of tumor enriched or depleted IPA genes in each cancer type. NA: not regulated.

**Table S3:** Functional enrichment of pan-cancer (A) tumor enriched and (B) depleted IPA genes (tumor vs. paired normal).

**Table S4:** (A) Summary of genes and samples truncated by truncating mutations or IPAs in human tumor tissues. TR: truncating mutations, DDR: DNA damage repair genes.

(B) List of genes and samples truncated by IPA, truncating mutations or both. TR: truncating mutations, DDR: DNA damage repair genes, TSG: tumor suppressor genes.

(C) Number of samples containing DDR genes that are truncated by IPA, TR or both. TR: truncating mutations.

(D) Number of samples carrying TSGs that are truncated by IPA, TR or both. TR: truncating mutations.

(E) List of tumor-enriched genes was associated with tumor mutation burdens.

**Table S5:** Number of IPA events positively or negatively correlated with polyadenylation/splicing factors.

**Table S6:** IPA-ncRNA isoforms positively correlated with signaling pathway members.

**Table S7:** (A) Number of genes, IPA and/or gene expression of which is associated with patient overall survival (OS).

(B) List of IPA and/or gene expression of which is associated with patient overall survival (OS). Consistency: Yes: both positively or negatively; No: one positively and another negatively; NA: one is missing

(C) List of IPA associated with immune profile.

(D) List of cancer types with subtype, stage, sex or race information.

(E) List of pan-cancer tumor-enriched IPA genes associated with patients’ cancer subtype, stage, sex or race.

